# Identification of candidate master transcription factors within enhancer-centric transcriptional regulatory networks

**DOI:** 10.1101/345413

**Authors:** Alexander J. Federation, Donald R. Polaski, Christopher J. Ott, Angela Fan, Charles Y. Lin, James E. Bradner

## Abstract

Regulation of gene expression through binding of transcription factors (TFs) to cis-regulatory elements is highly complex in mammalian cells. Genome-wide measurement technologies provide new means to understand this regulation, and models of TF regulatory networks have been built with the goal of identifying critical factors. Here, we report a network model of transcriptional regulation between TFs constructed by integrating genomewide identification of active enhancers and regions of focal DNA accessibility. Network topology is confirmed by published TF ChIP-seq data. By considering multiple methods of TF prioritization following network construction, we identify master TFs in well-studied cell types, and these networks provide better prioritization than networks only considering promoter-proximal accessibility peaks. Comparisons between networks from similar cell types show stable connectivity of most TFs, while master regulator TFs show dramatic changes in connectivity and centrality. Applying this method to study chronic lymphocytic leukemia, we prioritized several network TFs amenable to pharmacological perturbation and show that compounds targeting these TFs show comparable efficacy in CLL cell lines to FDA-approved therapies. The construction of transcriptional regulatory network (TRN) models can predict the interactions between individual TFs and predict critical TFs for development or disease.

## Introduction

Regulation of gene expression is achieved through cell-type specific interpretation of regulatory DNA. This regulatory DNA is bound by transcription factors (TFs) that recognize specific DNA sequence motifs and contain effector domains that regulate the proximal assembly and elongation of the transcriptional machinery [1]. Combinatorial TF regulation is the basis for a complex transcriptional regulatory network (TRN) within the cell capable of maintaining homeostasis, directing differentiation, and responding to extracellular stimuli [2,3]. Modeling these networks in simple organisms has provided insight into the evolutionary basis for network logic and allows testable predictions to be made on a controlled biological system [4].

While the complexity of mammalian TRNs has prevented the assembly of predictive systems-level models, many approaches have been applied to interrogate various components of these systems. Notable approaches include systematic identification of protein-protein interaction networks [5], gene co-expression networks [6], protein-gene interaction networks [7], master regulator identification [8,9], 3-dimsensional conformational networks [10], and expression perturbation networks [11]. A more direct approach for predicting TRNs specifically considers the physical interaction of TFs with DNA, as measured by heightened accessibility to the DNAseI enzyme [12]. However, these published algorithms only consider promoter-proximal DNaseI hypersensitive sites (DHSs), which exclude many distal regulatory elements from the analysis. Recent large-scale epigenome analyses have shown that cell-type specific regulatory activity primarily occurs in large enhancer elements distal from gene promoters [13,14], which are be captured in promoter-centric TRN models. Moreover, in addition to active enhancers, accessibility measurements identify insulator CTCF loci and poised regulatory elements [15,16], confounding TRN models of active, steady-state transcription.

Here we present a methodology for the assembly of TRNs inferred from DNA accessibility data within large enhancers defined by asymmetrically large histone acetylation (H3K27ac) domains. Integrating H3K27ac enhancer maps with DNA accessibility allows identification of discrete pockets containing regulatory operator sequences within large enhancers, improving on models of TRNs that only consider DNA accessibility. These TRNs accurately predict TF-enhancer interactions, identify known master regulatory TFs in model cell types, and can predict critical effector TFs in cancer for further pharmacological validation.

## Results and Discussion

We utilized publicly available ENCODE data from two human cell lines to validate our methods to construct enhancer-linked models of transcriptional regulatory networks: H1 human embryonic stem cells (hESCs) and the GM12878 lymphoblastoid cell line [17]. The strategy for network construction is summarized in Figure 1A. First, to allow network construction in the absence of transcriptome data, H3K27ac ChIP-seq signal at promoters is used as a surrogate to define the set of expressed genes in the cell. Promoter H3K27ac signal is highly correlated with RNA-seq and greater than 91% of genes predicted to be expressed by H3K27ac are confirmed by RNA-seq (Supplemental Figure 1). Should transcriptome data be available, these measurements can be used instead.

**Figure 1:**
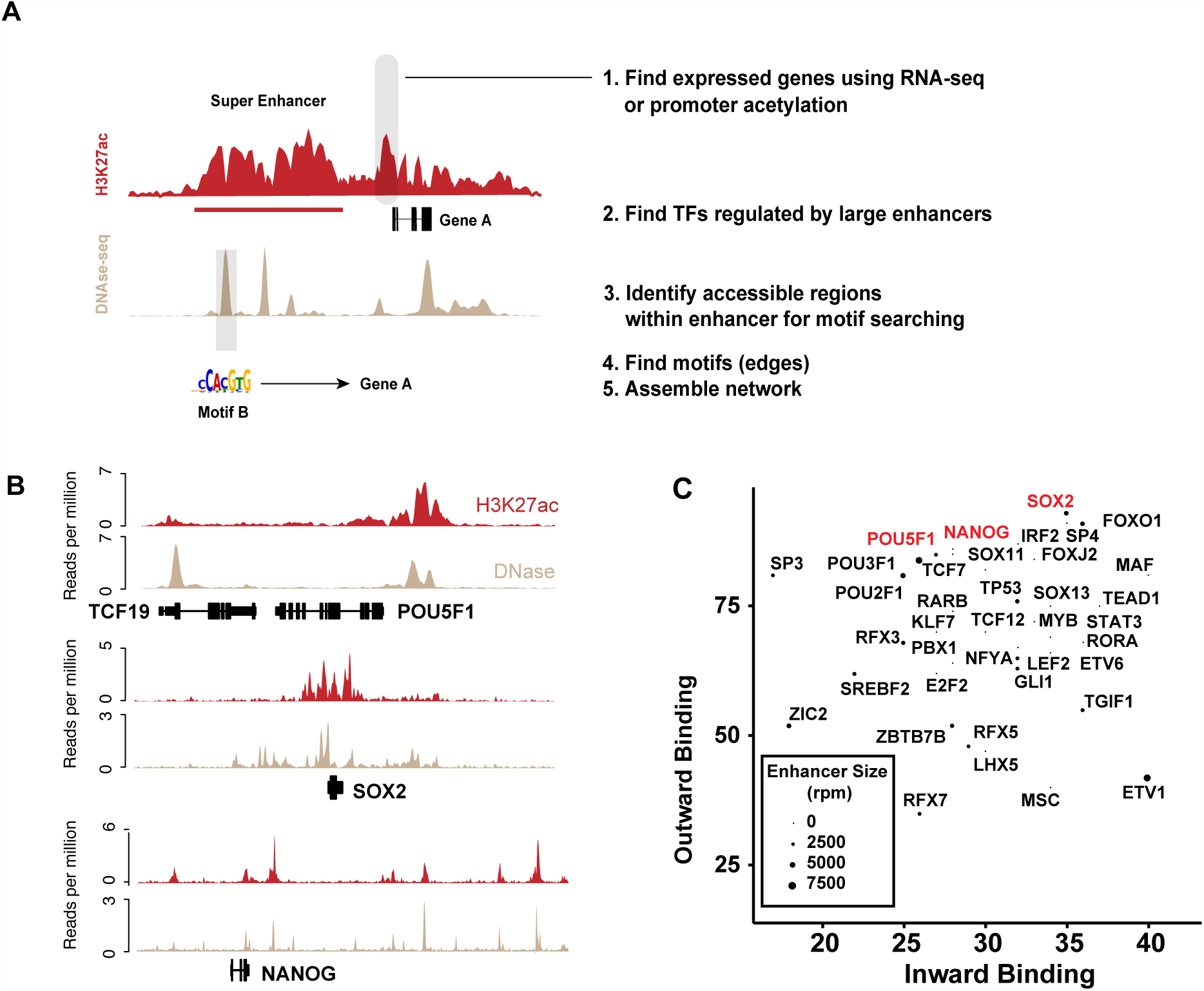
Integration of chromatin accessibility and ChIP-seq to assemble enhancer-linked transcriptional networks. (A) A schematic representing the Coltron algorithm for construction of TRNs. (B). Gene tracks showing super enhancers regulating the OSN master TFs in H1 cells. (C) In-degree vs. Out-degree for TFs within the H1 TRN.

Next, enhancers identified from focal H3K27ac peak signal are stitched and ranked by ChIP-seq read density using the rank ordering of super enhancers (ROSE2) algorithm [14,18]. Large enhancers have a disproportionately large share of cell-type specific regulatory DNA [14, 19], so these were used as the basis for TRN construction in this study. Within large H3K27ac enhancer domains, genome accessibility measurements are used to identify focal regions for motif finding. For network validation, chromatin accessibility was measured with DNAse-seq. ATAC-seq, which measures accessible chromatin through the activity of the Tn5 DNA transposase enzyme [20], was also considered as a substitute for DNAse-seq for edge prediction. Both measurements are highly correlated and result in networks with high degrees of similarity (95% shared edges, Supplemental Figure 2). DNA sequence underlying the relevant accessibility peaks are extracted used in the find individual motif occurrences (FIMO) algorithm to identify known TF operator sequences [21]. An edge is drawn between the TF and its target when a motif is discovered in the enhancer assigned to the target gene.

As expected in hESCs, the three master regulatory factors [22]- POU5F1 (also known as OCT4), SOX2 and NANOG (OSN) are regulated by super enhancers (Figure 1B), predicted to bind their own super enhancers, and bind super enhancers of the other master factors [13]. The method predicts that every node in the TRN is bound by at least one of these factors, consistent with their central role in pluripotency (Supplementary Figure 3). Plotting the in-degree (number of TFs binding the enhancer of a node TF) and out-degree (number of TF enhancers bound by a node TF) of each node in two dimensions allows for the visualization of the overall connectivity of nodes in the network (Figure 1C). Genes with known roles in pluripotency, FOXO1 [23], RARB [24], STAT3 [25], MYB [26] and OSN show remarkably high connectivity (top 10% of all TFs in the network). Similar results are seen for GM12878 (Supplementary Figure 4), with known lymphoid master regulators and IRF4, RUNX3 and SPI1 highly connected in the transcriptional network [6,27-29].

We then utilized TF ChIP-seq, when available for relevant TFs, to validate edges in the network. Predicted targets of OSN were found to contain ChIP-seq peaks at the sites predicted by the TRN (Supplementary Figure 5). Our predictions for NANOG and SOX2 edges performed with 87% and 83% accuracy, respectively. The only available dataset for POU5F1 identified fewer significantly enriched peaks (<50% of NANOG peaks), but still confirmed 75% of our predictions. False negative rates ranged from 2-4% in H1 hESCs. TFs in the GM12878 network performed at better rates, with accuracy ranging from 82-100%, possibly owing to the availability of higher quality TF ChIP-seq.

Using this method, we then constructed TRNs for all cell types with available high-quality data from the Roadmap Epigenomics Project [30]. With the goal of identifying metrics to prioritize master TFs for functional follow-up, we studied three different strategies to prioritize TFs in TRNs from well-characterized cells in the hematopoietic lineage (CD34+ hematopoietic stem cells, CD3+ T cells, and CD20+ B cells). First, common network centrality calculations were considered and used to rank each TF in a TRN based on the its connectivity within the network. We tested 6 published algorithms to define betweenness [31], closeness [32], total-degree, eigenvector [33], in-degree, and out-degree centrality. In general, ranking TFs by the published centrality scores resulted in highly correlated outcomes (Pairwise Spearman correlations > 0.8, Figure 2A), The exception to this pattern is in-degree centrality, which correlates poorly with the other metrics and performs worst at TF prioritization. Master TFs defined in the literature for hESCs and different hematopoietic tissues were ranked highly by all centrality metrics except for in-degree centrality (Supplementary Figure 6). In all tissues examined, TRNs built with this method outperformed previously published promoter-proximal TRNs at identifying master TFs with high centrality measurements (Supplementary Figure 7).

**Figure 2.**
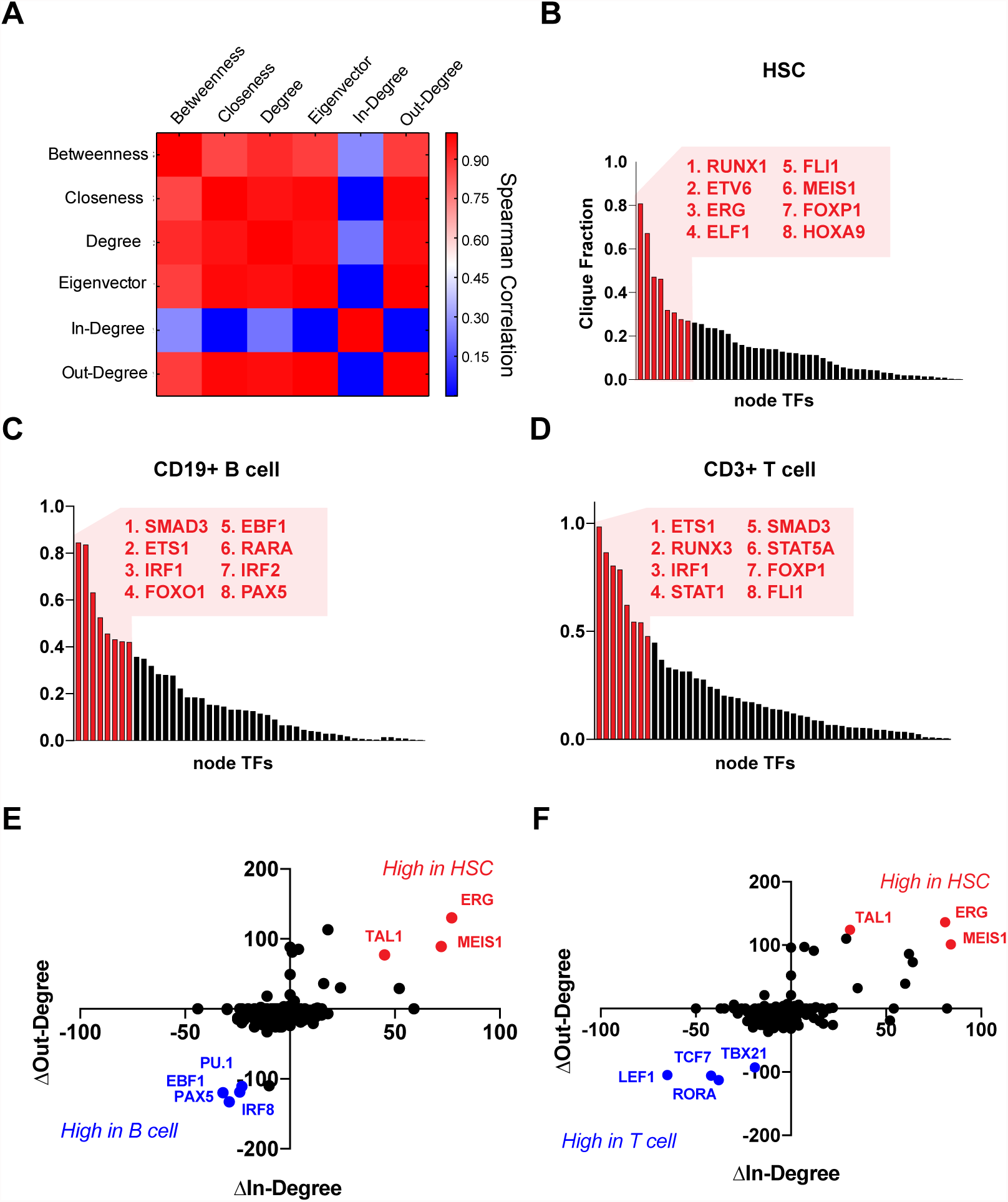
Comparison of network centrality calculations for periodization of master regulatory TFs. (A) Correlation matrix comparing TF rankings as calculated by each indicated metric. TFs ranked by clique fraction in (B) CD34+ HSCs, (C) CD19+ B cells and (D) CD3+ T cells. (E, F) Scatterplot showing changes in In-degree and Out-degree when comparing HSC vs B cell and HSC vs T cell.

While these centrality metrics consider a TF’s local interactions within the TRN, we sought to devise a metric that captured a TF’s expanded role within fully interconnected self-reinforcing network motifs (also referred to as cliques). These network motifs are common between sets of master regulator transcription factors and are well-described in the context of pluripotency maintenance hESCs [22], where OSN share a clique. We searched for cliques within the Roadmap TRNs and found that they occur frequently, with an average of 64 cliques per sample. When the TFs participating in these cliques were studied, we found a small set of TFs dominate the membership of cliques globally within the TRN. To quantify this, we defined ‘clique fraction’ for a given TF as the number of cliques for which the TF is a member divided by total cliques. CD34+ HSCs show many master regulators of hematopoiesis with highly ranked clique fractions including ERG, FLI1 and HOXA9 (Figure 2B) [27]. Similarly, the B cell and T cell TRNs clique-fraction rankings also prioritize many TFs with roles in B cell and T cell biology, including some TFs with general roles in hematopoiesis (SMAD3, ETS1 and IRF1) and some lineage-specific master regulators (PAX5 in B cells or RUNX3 in T cells) [34].

Lastly, we considered an alternative strategy to identify nodes critical in TRNs by leveraging direct comparison of networks derived from two different biological conditions. As expected for similar cell types, when TRNs from CD34+ HSCs are directly compared to B cells and T cells, most TFs within the network have similar connectivity profiles. However, a small number (3-5) of TFs are outliers and show large differentially connectivity between the two networks, and these TFs are well-studied regulators of that specific biology. TAL1, ERG and MEIS1 were highly connected in HSCs compared to both T cells and B cells, and each of these factors has a well-established function in HSC maintenance and function [27]. The B cell master regulators PAX5, EBF1 and PU.1 are hyperconnected in B cells compared to HSCs while T cell master factors LEF1, RORA, TCF7 and TBX21 are hyperconnected in the T cell TRNs.

With the ability to predict TFs that play a central role in transcriptional regulation for healthy tissues, we hypothesized that TRNs may allow prioritization of TFs for functional validation as novel cancer dependencies. To test this hypothesis in an unbiased manner, we integrated large scale RNAi experiments analyzed by DEMETER2 [35] with TRNs from cancer cell lines included in the Roadmap data [30]. Four cell lines were included in both datasets – K562, HepG2, A549 and HELA. Considering a stringent cutoff to define an essential gene (DEMETER2 score < 0.25) all cell lines except for HELA show an enrichment for essential genes within the top 50 TFs as ranked by clique fraction (Supplementary Figure 8). As explored in the hematopoietic samples, we expect even greater enrichments for essential genes if cancer TRNs are compared against TRNs from relevant normal tissues.

To generate a dataset to directly interrogate the essentiality of highly-connected TFs in a cancer context, we collected H3K27ac ChIP-seq and ATAC-seq data from the MEC1 chronic lymphocytic leukemia (CLL) cell line and closely related healthy CD19+ primary human B cells. TRN construction in MEC1 identified many previously studied TF oncogenes in CLL (Figure 3A) including MYC, ETS1, IRF4 [36-38]. Differential analysis of MEC1 vs B cells uncovered 4 TFs that are highly connected in CLL that can be pharmacologically targeted with known small molecule compounds – PPARA, RARA, IKZF1, NFATC1/2 (Figure 3B). All 4 TFs are highly ranked by centrality metrics and clique fraction (Figure 3C, Supplementary Figure 9). When MEC1 cells were treated with small molecule compounds targeting each TF, all compounds show significant effects on cell proliferation in dose response (Figure 3C). As a positive control, the FDA-approved CLL therapies Ibrutinib and Idelalisib were tested and show equal or less potent efficacy in MEC1 that our 4 TF-targeting compounds (Figure 3D).

**Figure 3:**
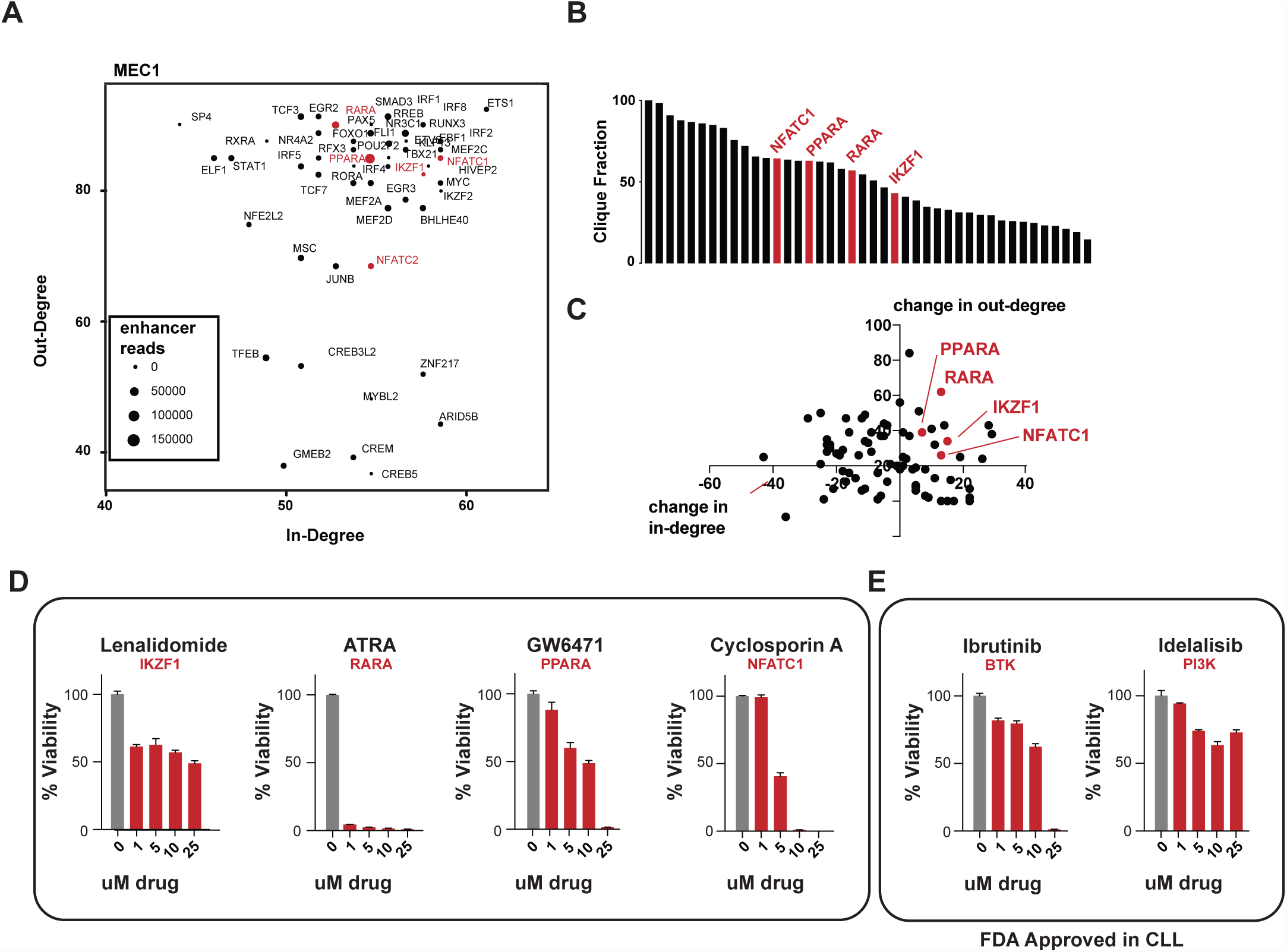
Pharmacological validation of CLL-specific TRN TFs. (A) In-degree vs Out-degree of the MEC1 CLL cell line (B) TF ranking by clique fraction in MEC1, highlighting the TFs targetable by small molecule compounds. (C) TRN comparison between MEC1 and CD19+ B cells, highlighting TFs targetable by small molecule compounds. (D, E) Survival curves of MEC1 cells treated with the indicated compounds in dose response after 24 hours.

In conclusion, we present a methodology for constructing models of TRNs connected through large enhancers that integrate DNA accessibility data for robust TF-binding predictions. The method requires only two genome-wide measurements, making it amenable for profiling primary tissue samples constrained by small sample inputs. Predicted TRN centrality measurements identify TFs with well-established biological roles in development and can identify novel cancer dependencies. When available, these dependencies can be targeted by FDA-approved small molecule compounds, providing an opportunity for drug repurposing in CLL.

The source code for TRN construction is freely available to the broader community for use and further development. Additionally, TRNs derived from all tissues profiled with high quality by the Roadmap Epigenomics Consortium have been assembled and are freely available. We expect these techniques to aid in the interpretation of large epigenomic datasets and provide a pathway to prioritize functional validation of TFs in developmental biology and therapeutic development.

## Methods

### Datasets and processing

Datasets were obtained through the publicly available ENCODE data repository [17] and the Roadmap Epigenomic Consortium data repository [30]. ChIP-seq data from both sources was processed using a standardized pipeline. Peak finding analysis was performed using Model-based analysis of ChIP-seq (MACS) version 1.4 with a p-value cutoff value 1E-6 [39]. Identified peak regions were then stitched together using the rank ordering of super enhancers (ROSE) algorithm [13]. A maximum linking distance of 12500 bp was used as the stitching parameter along with a TSS exclusion region of 1000 bp. Gene tracks were plotted with BamPlot (https://github.com/BradnerLab/).

### Software and dependencies

The TRN software uses the following dependencies:

- Bamliquidator – version 1.2.0
- Samtools – version 0.1.19
- FIMO – version 4.91
- NetworkX – version 1.8.1

### Transcriptional Regulatory Network

Detailed documentation and source code for TRN construction can be found at https://github.com/BradnerLab/. In brief, H3K27ac ChIP-seq levels are calculated at all promoters and the top 50% of promoters are considered ‘expressed’ as possible nodes for the TRN networks. Enhancer domains precalculated in ROSE are assigned to nearby expressed transcripts. Enhancer -associated transcription factors (TF) are then selected from the lists of enhancer-associated genes using a list of 2189 unique TF transcripts. Within enhancers assigned to TFs, DNAse or ATAC peaks are identified and the underlying sequence is extracted for motif searching. FIMO was used to identify motifs within these sequences using a p-value of 1e-4 for motifs taken from TRANSFAC and JASPAR. An edge is drawn from a TF to a target when its motif is found in the enhancer regulating the target and these edges are complied to form the complete TRN for further analysis.

### Centrality calculations

Centrality calculations were carried out using the NetworkX python module. Networks were analyzed using six centrality measures, in-degree, out-degree, total degree, betweenness, closeness, and eigenvector (Pavlopoulos *et al.* 2011). In-degree and out-degree centrality are calculated for each node by summing the number of incoming and outgoing adjacent edges, respectively. Total degree centrality is calculated by summing the total number of edges adjacent to the given vertex.

Betweenness centrality of a vertex is defined as the number of shortest paths between each pair of two vertices that pass through the given vertex. The equation for calculating the betweenness centrality of the vertex is:

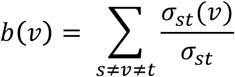

Where *σ_st_* is the total number of shortest paths between vertex s and vertex t, and *σ_st_*(*v*) is the number of those shortest paths that contain vertex v.

Closeness centrality of a vertex is defined as the reciprocal of the sum of the short distance between that vertex and every other vertex in the network. Closeness centrality was calculated using the following expression:
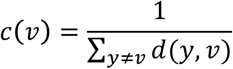

Where *d*(*y*, *v*) is the distance between the vertex of interest v and another vertex y.

Eigenvector centrality is calculated by solving the following eigenvalue equation:

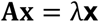

Where **A** is the adjacency matrix of the graph and λ is the greatest eigenvalue of **A**. Eigenvector centrality for a vertex v is defined as the v^th^ component of eigenvector associated with this eigenvalue.

### Comparative network analysis

The TRN methodology for network comparison is identical to the single network approach, with the following exceptions. The TF node set is comprised of the union of the node sets from both networks, assuming that TF is expressed. The enhancer set is comprised of the combined, merged enhancers from both networks.

### Cell culture

For cell line assays, cells were plated in 384-well plates at a seeding density of 5,000 cells/mL in RPMI media supplemented with 10% FCS. One day after seeding, cells were treated with compounds resuspended as DMSO stock solutions via pin-transfer robots (Janus Workstation, Perkin Elmer). Four days following treatment, 1/10 volume of AlamarBlue Cell Viability Reagent (Life Technologies) was added and incubated with live cells for 24 hours. After incubation, fluorescence was read on an Envision multi-well plate reader (PerkinElmer). Compounds were obtained from commercial sources (Selleck).

### ChIP-seq

Chromatin immunoprecipitation was performed with 3 cells per sample using an anti-H3K27ac specific antibody (Abcam #ab4729). ChIP procedure was performed as previously described [40] Samples were sequenced on a Illumina HiSeq 2000 or 2500, paired-end, 100 x 100 cycles. Cell lines were sequenced on on an Illumina NextSeq in single-end mode 75 cycles. Raw paired-end sequencing data was mapped to the hg19 build of the human genome with Bowtie2 with default settings and the parameters –p 4 –k 1 [41]. Mapped reads were filtered to remove duplicate reads, and to regions in the ENCODE blacklist. MACS was used for peak identification with a p-value cutoff of 1e-6.

### ATAC-seq

ATAC-seq was performed with 50,000 viable cells as described with minor modifications [42]. Transposition reactions were performed for 1 hour at 37° C, followed by purification and sample barcoding by PCR. Samples were sequenced on an Illumina HiSeq 2000 or 2500 in paired-end mode with 100 x 100 cycles. ATAC-seq for MEC1, MEC2, OSU-CLL, and Cll cells was performed as above after cells were grown in RPMI-1640 supplemented with 10% FCS. ATAC-seq libraries were sequenced on an Illumina NextSeq in paired-end mode with 75 x 75 cycles. Raw paired-end sequencing data was mapped to the hg19 build of the human genome with Bowtie2 with default settings and the parameters –p 4 –k 1. Mapped reads were filtered to remove duplicate reads, those reads mapping to chromosome M and to regions in the ENCODE blacklist. Reads mapping to the forward strand were shifted forward 4bp, and reads mapping to the reverse strand were shifted backwards 5bp. ZINBA was then used to find peaks with extension=200, winSize=300 and offset=75 and default settings [43].

## Supplementary Figure Legends

**Supplementary Figure 1:**
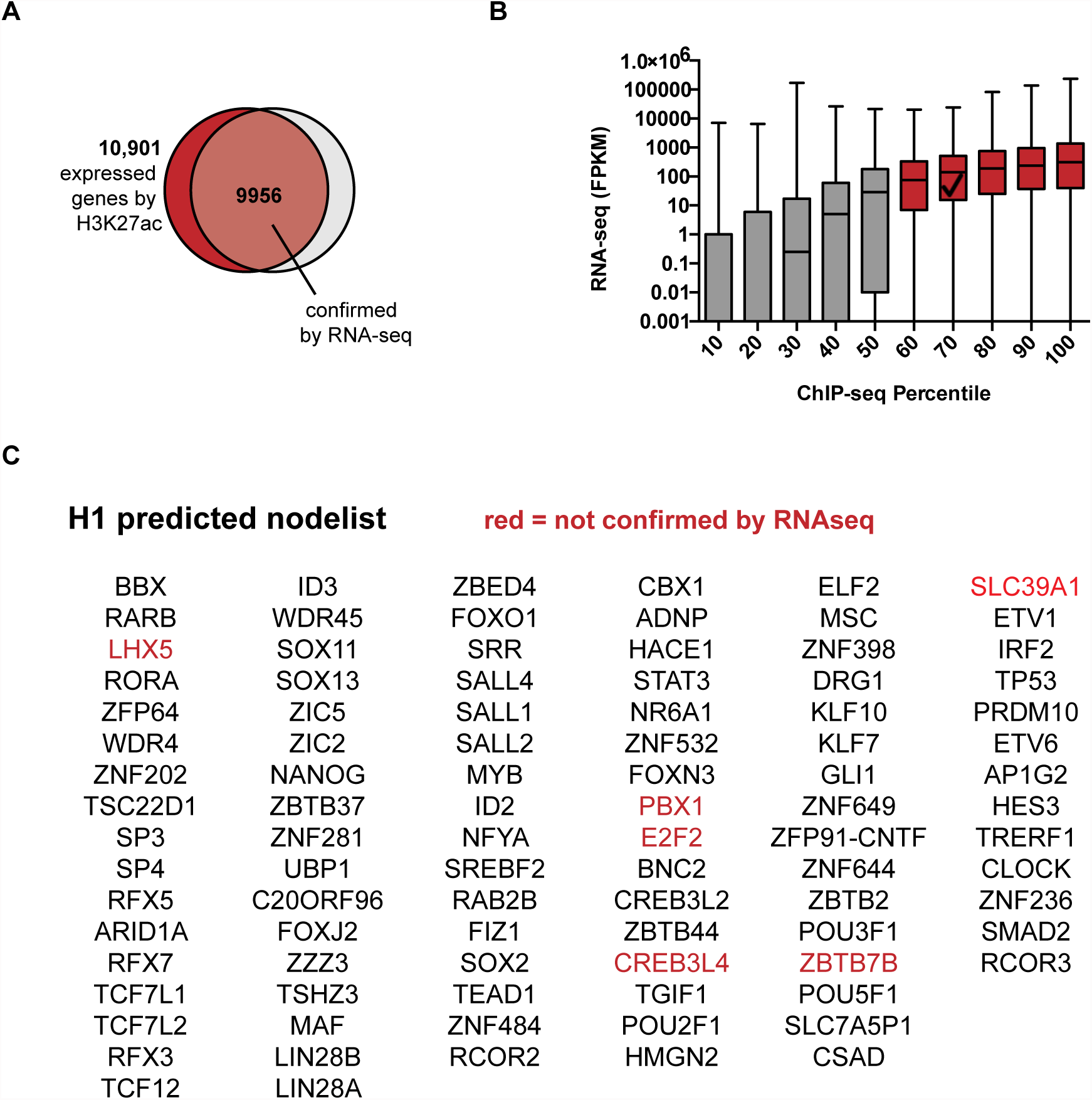
Promoter H3K27ac as a surrogate for RNA expression levels. All genes (hg19) were ranked by RNA-seq and H3K27ac promoter signal (+/- 1000 bp from TSS) in H1 cells. (A) Overlap of genes with FKPM > 1.0 and genes in the top 50% of promoter acetylation signal. (B) RNA-seq distributions for genes across deciles of H3K27ac promoter acetylation. (C) TF genes predicted as active for TRN construction in H1 cells by H3K37ac promoter signal, genes in red show RNA-seq FKPM < 1.0.

**Supplementary Figure 2:**
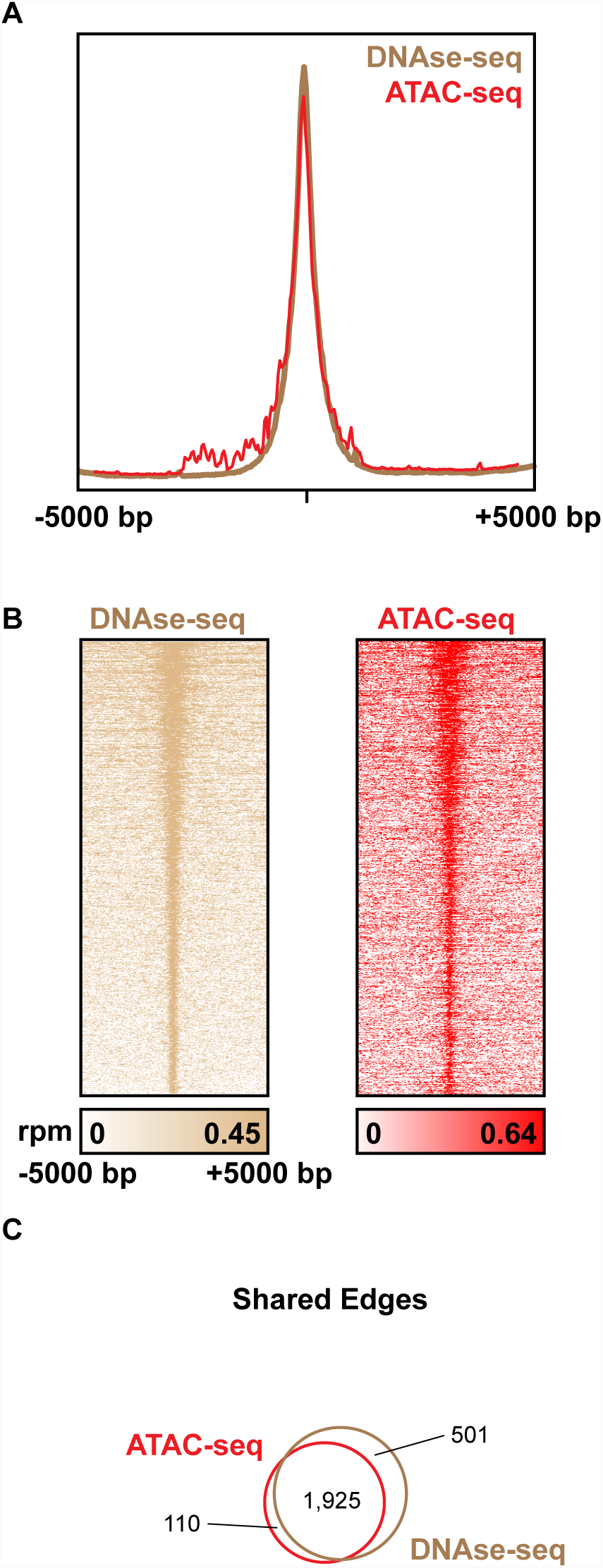
Comparison of DNase-seq and ATAC-seq as accessibility measurements for TRN construction. H1 TRNs were constructed using DNA accessibility measurements using published DNase-seq and ATAC-seq data. (A) Meta-track showing significant overlap of read density of DNase-seq and ATAC-seq at all DHSs (B) Read density at all DHSs ranked by total DNase-seq read density. (C) Overlap of network edges between TRNs constructed from each accessibility measurement.

**Supplementary Figure 3:**
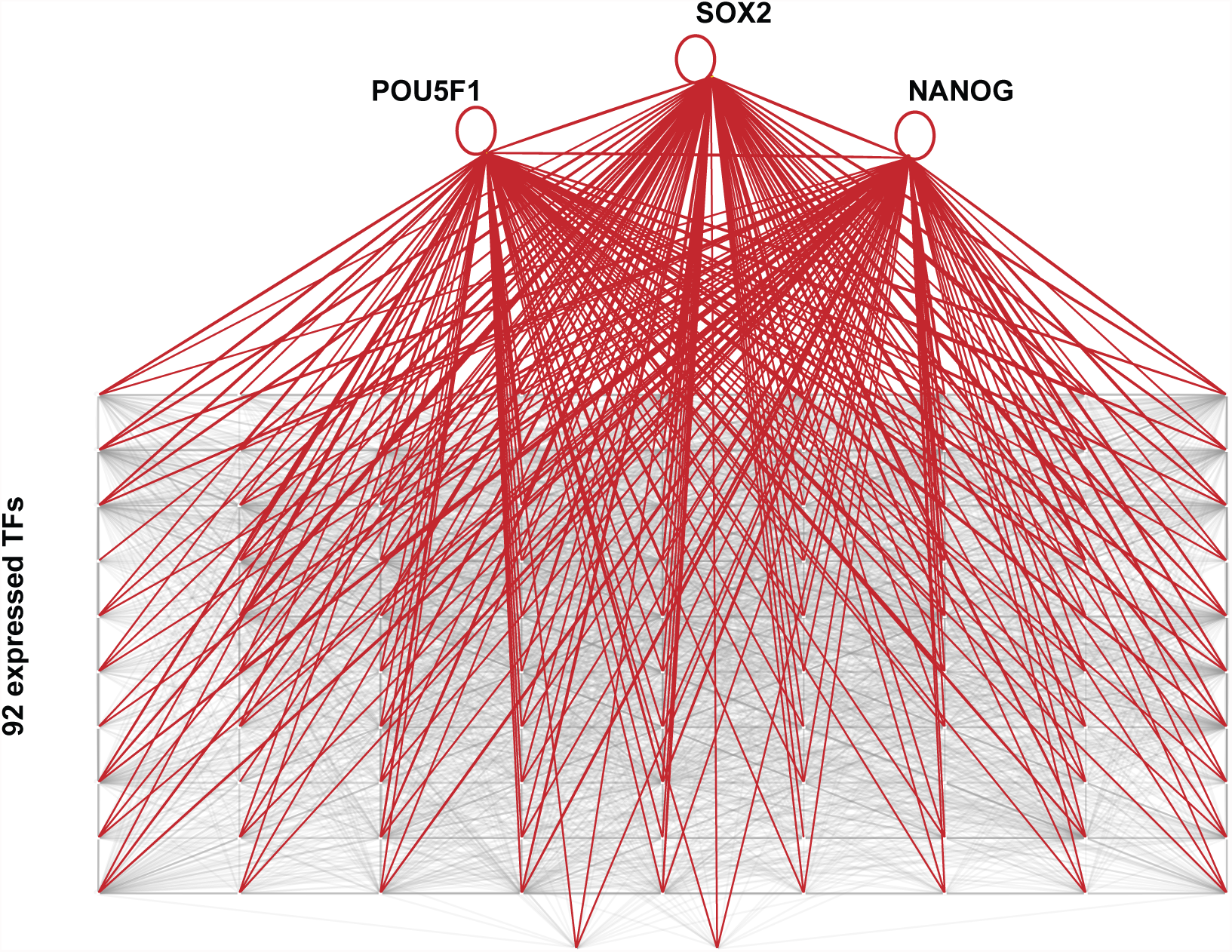
The H1 super-enhancer centric TRN. All nodes in the H1 TRN. Edges between the master regulators OSN and other node TFs are highlighted in red.

**Supplementary Figure 4:**
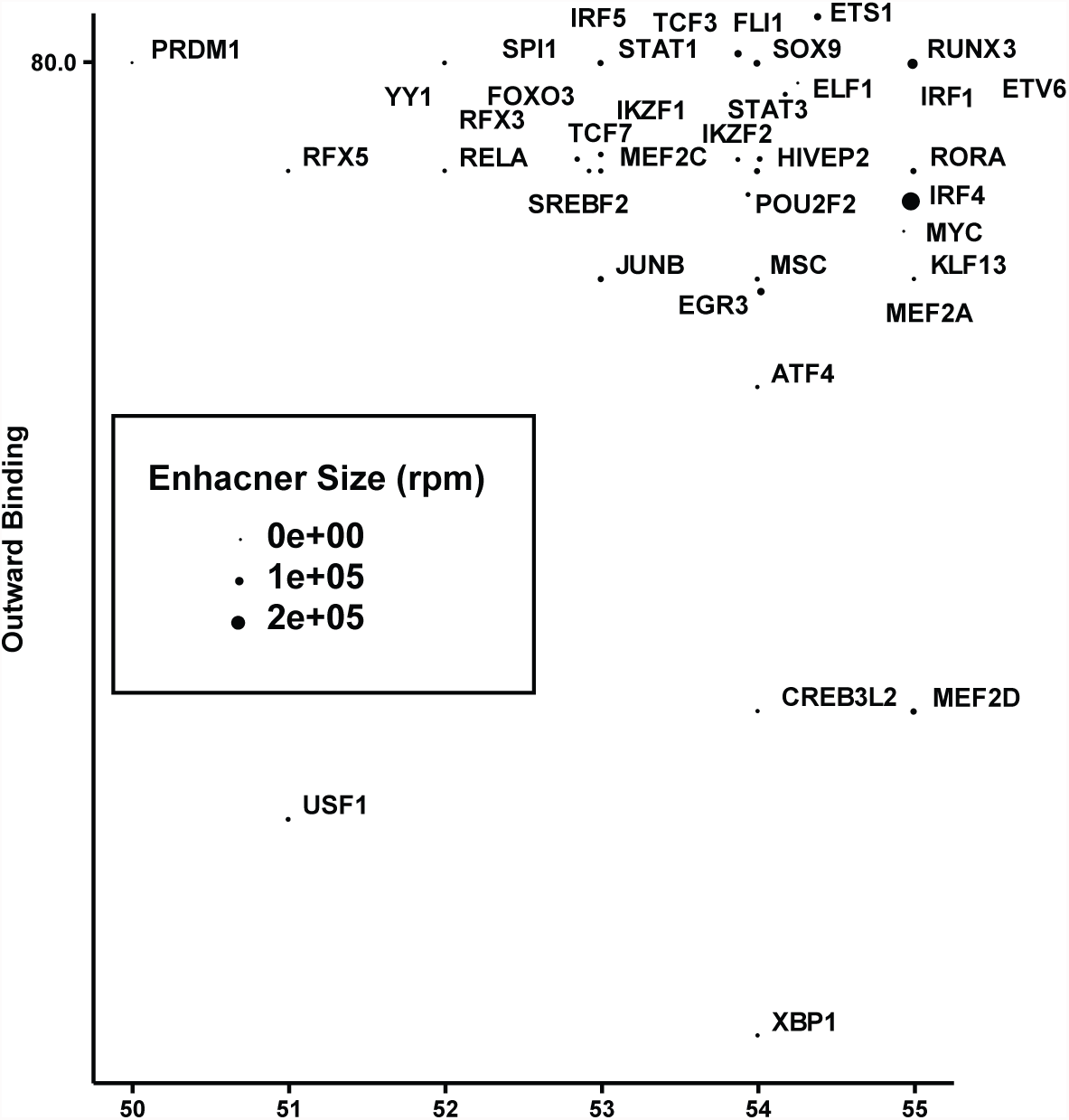
The GM12878 super-enhancer centric TRN. In-degree vs Out-degree in the TRN of GM12878 lymphoblast cells.

**Supplementary Figure 5:**
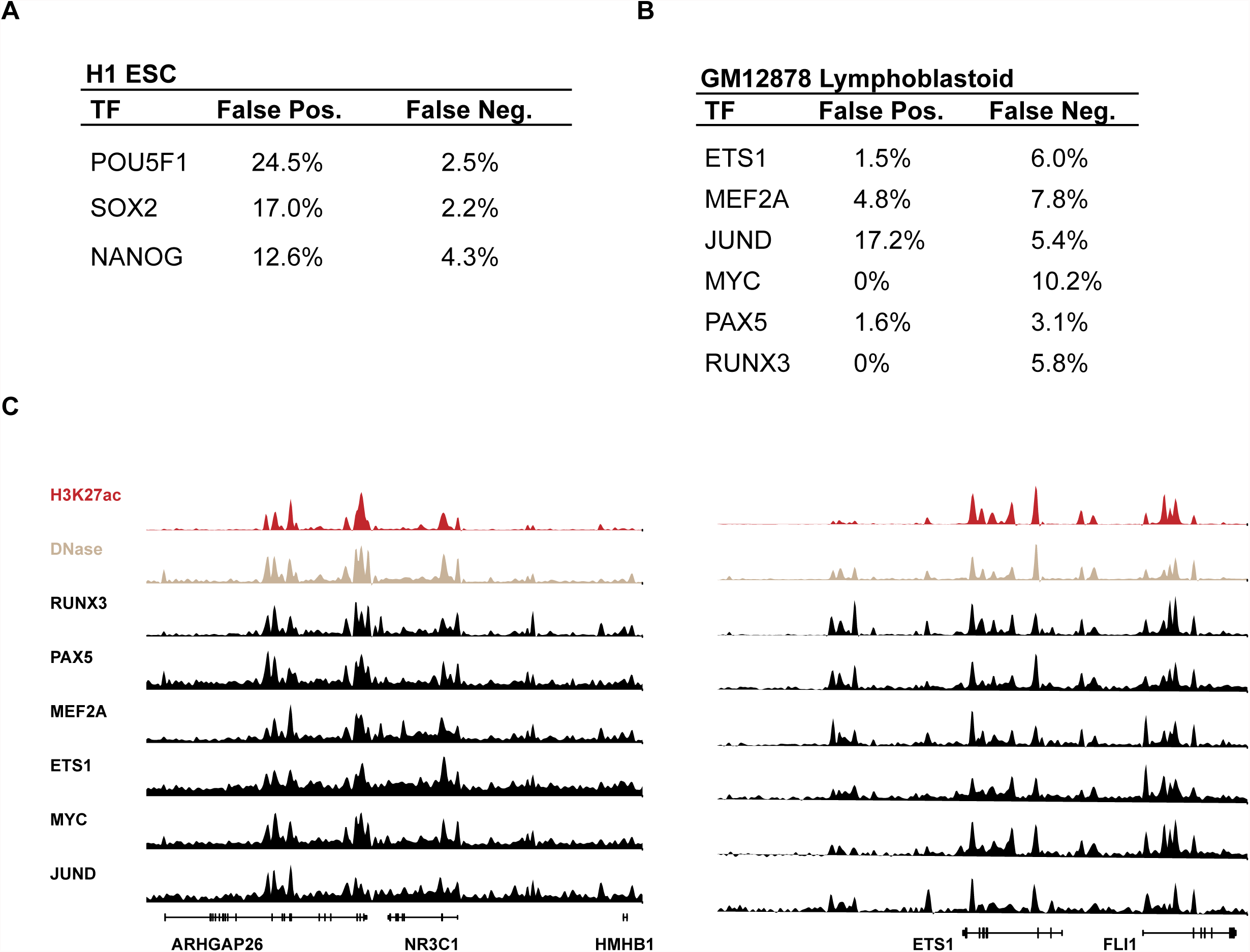
Edge validation in H1 and GM12878 TRNs using ChIP-seq data. False-positive and false-negative rates for TFs in TRNs using publicly available ChIP-seq data at ‘true positive’ edges in (A) H1 cells and (B) GM12878 cells. (C) Gene tracks highlighting large enhancer regions with TF binding validated by ChIP-seq

**Supplementary Figure 6:**
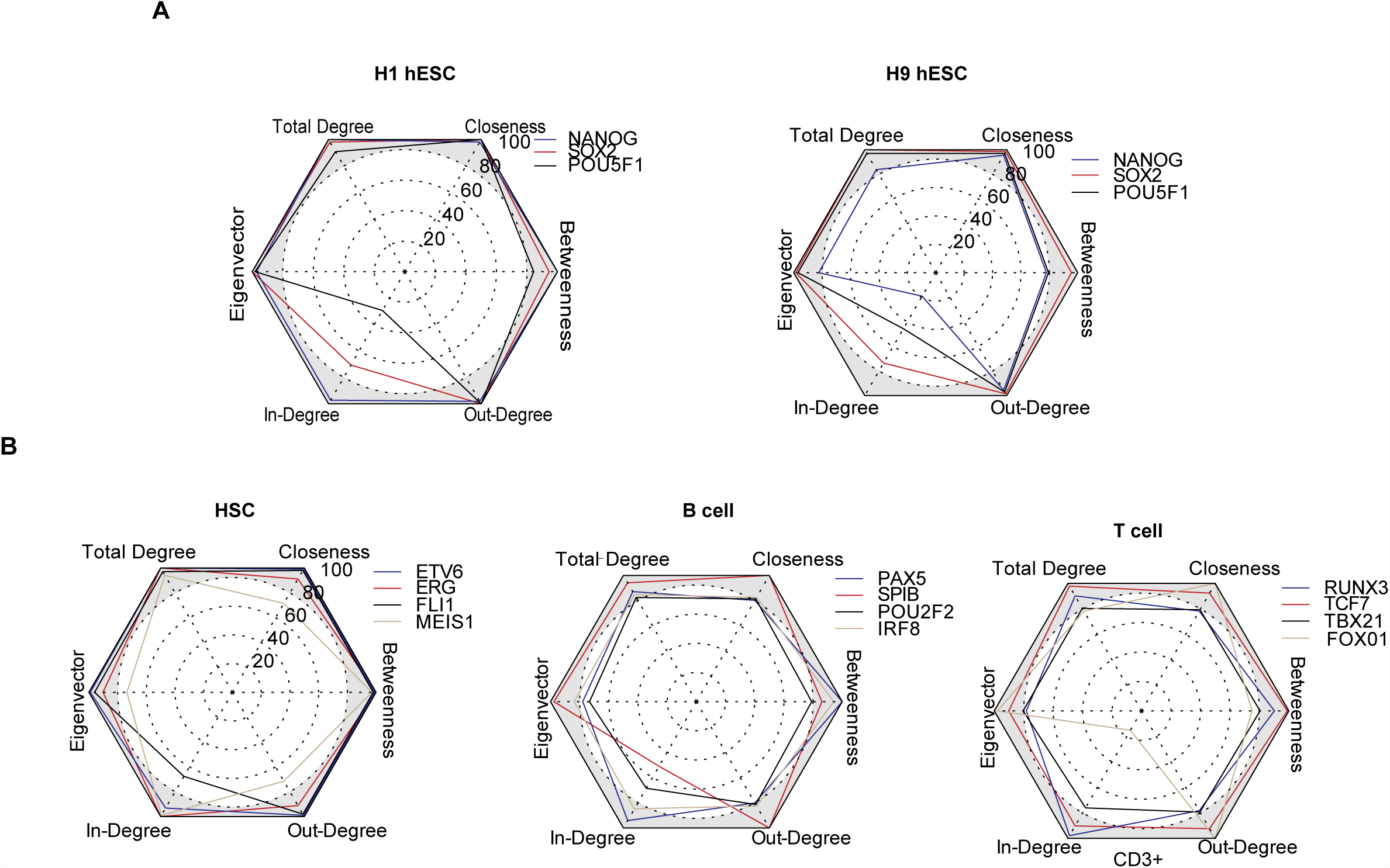
Comparison of 6 centrality metrics for priority ranking of TFs. For each cell type, rankings of master TFs defined in the literature are shown for each centrality measurement. The grey-shaded region represents a ranking in the top 10% of TFs. (A) H1 and H9 hESC cells (B) Roadmap cells from hematopoiesis – CD34 HSCs, CD3 T cells and CD19 B cells.

**Supplementary Figure 7:**
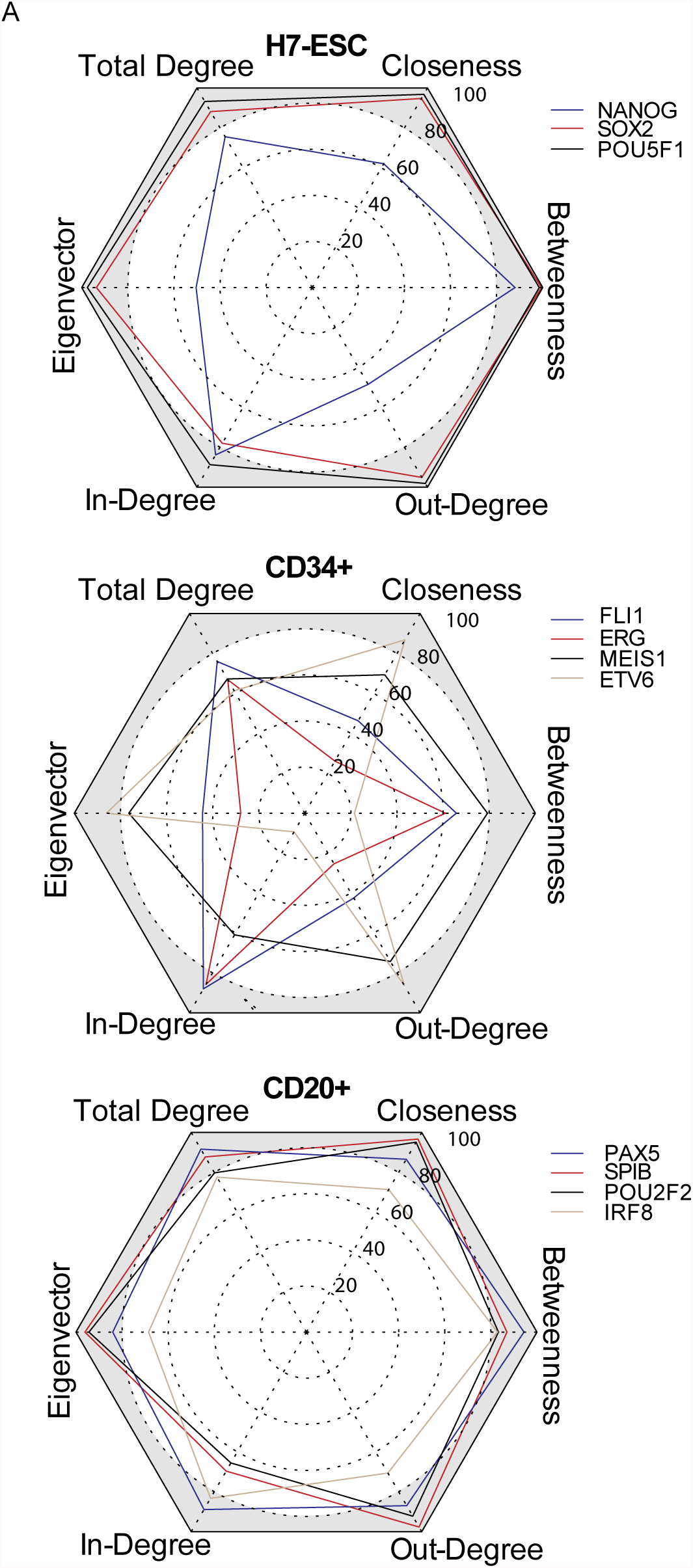
Comparison of 6 centrality metrics for priority ranking of TFs from networks published in Neph et al [**12**]. For each cell type, rankings of master TFs defined in the literature are shown for each centrality measurement.

**Supplementary Figure 8:**
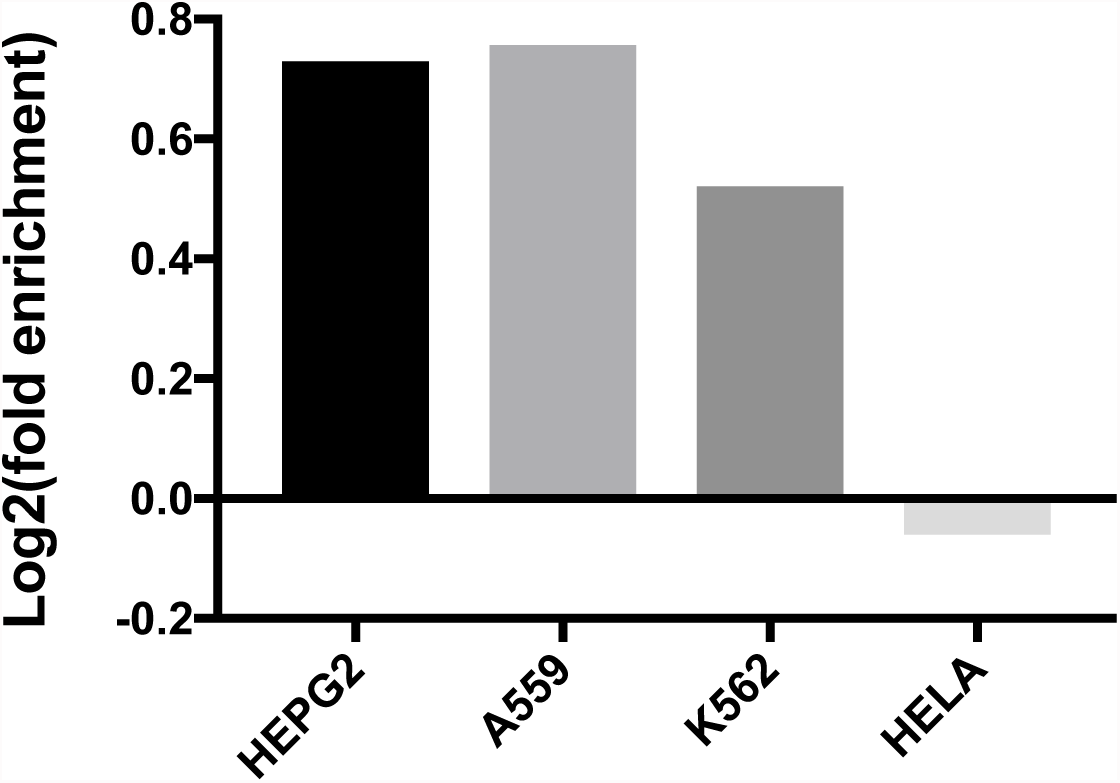
Enrichment scores for likelihood of finding essential genes within top TRN TFs. For each cell line, TRNs were constructed and TFs were ranked by clique fraction. For the top 50 TFs as ranked by clique fraction, the rate of identifying an essential gene was calculated and enrichment was calculated by comparing to the likelihood of finding an essential gene when considering all expressed genes.

## References

1. Lambert, S. A. et al. The Human Transcription Factors. Cell 172, 650–665 (2018).

2. Alon, U. Network motifs: theory and experimental approaches. Nature Reviews Genetics 8, 450–461 (2007).

3. Davidson, E. Emerging properties of animal gene regulatory networks. Nature 468, 911–920 (2010).

4. Karr, J. R. et al. A whole-cell computational model predicts phenotype from genotype. Cell 150, 389–401 (2012).

5. Rolland, T. et al. A Proteome-Scale Map of the Human Interactome Network. Cell 159, (2014).

6. Novershtern, N. et al. Densely interconnected transcriptional circuits control cell states in human hematopoiesis. Cell 144, 296–309 (2011).

7. Bass, J. et al. Human Gene-Centered Transcription Factor Networks for Enhancers and Disease Variants. Cell 161, 661–73 (2015).

8. Della Gatta, G. et al. Reverse engineering of TLX oncogenic transcriptional networks identifies RUNX1 as tumor suppressor in T-ALL. Nat. Med. 18, 436–40 (2012).

9. Carro, M. S. et al. The transcriptional network for mesenchymal transformation of brain tumours. Nature 463, 318–25 (2010).

10. Sandhu, K. et al. Large-Scale Functional Organization of Long-Range Chromatin Interaction Networks. Cell Reports 2, 1207–1219 (2012).

11. Yosef, N. et al. Dynamic regulatory network controlling TH17 cell differentiation. Nature (2013). doi:10.1038/nature11981

12. Neph, S. et al. Circuitry and dynamics of human transcription factor regulatory networks. Cell 150, 1274–86 (2012).

13. Hnisz, D. et al. Super-enhancers in the control of cell identity and disease. Cell 155, 934–47 (2013).

14. Whyte, W. A. et al. Master transcription factors and mediator establish super-enhancers at key cell identity genes. Cell 153, 307–19 (2013).

15. Liu, M. et al. Genomic discovery of potent chromatin insulators for human gene therapy. Nat. Biotechnol. 33, 198–203 (2015).

16. Zentner, G., Tesar, P. & Scacheri, P. Epigenetic signatures distinguish multiple classes of enhancers with distinct cellular functions. Genome Research 21, 1273–1283 (2011).

17. Dunham, I. et al. An integrated encyclopedia of DNA elements in the human genome. Nature (2012). doi:10.1038/nature11247

18. Lovén, J. et al. Selective inhibition of tumor oncogenes by disruption of super-enhancers. Cell 153, 320–34(2013).

19. Dowen, J. et al. Control of Cell Identity Genes Occurs in Insulated Neighborhoods in Mammalian Chromosomes. Cell (2014). doi:10.1016/j.cell.2014.09.030

20. Buenrostro, J. D., Giresi, P. G., Zaba, L. C., Chang, H. Y. & Greenleaf, W. J. Transposition of native chromatin for fast and sensitive epigenomic profiling of open chromatin, DNA-binding proteins and nucleosome position. Nat. Methods 10, 1213–8 (2013).

21. Charles E. Grant, Timothy L. Bailey and William Stafford Noble, “FIMO: Scanning for occurrences of a given motif”, Bioinformatics 27(7):1017–1018, (2011).

22. Boyer, L. A. et al. Core transcriptional regulatory circuitry in human embryonic stem cells. Cell 122, 947–56(2005).

23. Zhang, X. et al. FOXO1 is an essential regulator of pluripotency in human embryonic stem cells. Nat. Cell Biol. 13, 1092–9 (2011).

24. Wang, W. et al. Rapid and efficient reprogramming of somatic cells to induced pluripotent stem cells by retinoic acid receptor gamma and liver receptor homolog 1. Proceedings of the National Academy of Sciences (2011). doi:10.1073/pnas.1100893108

25. Raz, R., Lee, C. K., Cannizzaro, L. A., d’ Eustachio, P. & Levy, D. E. Essential role of STAT3 for embryonic stem cell pluripotency. Proc. Natl. Acad. Sci. U.S.A. 96, 2846–51 (1999).

26. Zhan, M. et al. The B-MYB transcriptional network guides cell cycle progression and fate decisions to sustain self-renewal and the identity of pluripotent stem cells. PLoS ONE 7, e42350 (2012).

27. Moignard, V. et al. Characterization of transcriptional networks in blood stem and progenitor cells using high-throughput single-cell gene expression analysis. Nature Cell Biology (2013). doi:10.1038/ncb2709

28. Whiteman, H. J. & Farrell, P. J. RUNX expression and function in human B cells. Crit. Rev. Eukaryot. Gene Expr. 16, 31–44 (2006).

29. Carotta, S. et al. The transcription factors IRF8 and PU.1 negatively regulate plasma cell differentiation. Journal of Experimental Medicine (2014). doi:10.1084/jem.20140425

30. Consortium, R. E. et al. Integrative analysis of 111 reference human epigenomes. Nature 518, 317 (2015).

31. Brandes, U. A faster algorithm for betweenness centrality*. The Journal of Mathematical Sociology 25, 163–177(2001).

32. Freeman, L. C. Centrality in social networks conceptual clarification. Social Networks 1, 215–239 (1978).

33. Pavlopoulos, G. et al. Using graph theory to analyze biological networks. BioData Mining (2011). doi:10.1186/1756-0381-4-10

34. Cobaleda, C., Jochum, W. & Busslinger, M. Conversion of mature B cells into T cells by dedifferentiation to uncommitted progenitors. Nature 449, 473–477 (2007).

35. 1McFarland, J. M. et al. Improved estimation of cancer dependencies from large-scale RNAi screens using model-based normalization and data integration. (2018). doi:10.1101/305656

36. Huh, Y. O. et al. MYC translocation in chronic lymphocytic leukaemia is associated with increased prolymphocytes and a poor prognosis. Br J Haematol 142, 36–44 (2008).

37. Shukla, V., Ma, S., Hardy, R. R., Joshi, S. S. & Lu, R. A role for IRF4 in the development of CLL. Blood 122, 2848–2855(2013).

38. Fabbri, G. et al. Genetic lesions associated with chronic lymphocytic leukemia transformation to Richter syndrome. The Journal of Experimental Medicine 210, 2273–2288 (2013).

39. Zhang, Y. et al. Model-based Analysis of ChIP-Seq (MACS). Genome Biology (2008). doi:10.1186/gb2008-9-9-r137

40. Chapuy, B. et al. Discovery and Characterization of Super-Enhancer Associated Dependencies in Diffuse Large B-Cell Lymphoma. Cancer cell 24, 777–790 (2013).

41. Langmead, B. & Salzberg, S. L. Fast gapped-read alignment with Bowtie 2. Nature Methods 9, 357–359 (2012).

42. Buenrostro, J., Wu, B., Chang, H. & Greenleaf, W. ATAC-seq: A Method for Assaying Chromatin Accessibility Genome-Wide. Current protocols in molecular biology/ edited by Frederick M. Ausubel … [et al.] 109, 21.29.1–21.29.9 (2015).

43. Rashid, N. U., Giresi, P. G., Ibrahim, J. G., Sun, W. & Lieb, J. D. ZINBA integrates local covariates with DNA-seq data to identify broad and narrow regions of enrichment, even within amplified genomic regions. Genome Biology 12, R67 (2011).

